# Hierarchical Phase-Contrast Tomography Imaging: Applicability in biomedical research

**DOI:** 10.64898/2026.01.15.699614

**Authors:** Daniel Docter, Joseph Brunet, Vaishnavi Sabarigirivasan, Giel Lemmens, Birger Tielemans, Jon Sporring, Hector Dejea, Theresa Urban, Joanna Purzycka, Ramon Gorter, Judith Huirne, Veerle Michels, Jermo Hanemaaijer-van der Veer, Alexandre Bellier, David Stansby, Maximilian Ackermann, Roman Buelow, Danny Jonigk, Joseph Jacob, Jaco Hagoort, Stijn Verleden, Andrew C. Cook, Claire L. Walsh, Paul Tafforeau, Peter D. Lee, Bernadette S. de Bakker

## Abstract

**Objectives:** Hierarchical Phase-Contrast Tomography (HiP-CT) enables non-destructive, multi-scale imaging of whole human organs. We describe how HiP-CT is utilized for biomedical research within the Human Organ Atlas Hub through three case studies: mapping the enteric nervous system (ENS) of the human colon, analysing myocardial and AV conduction architecture in Tetralogy of Fallot (TOF), and characterizing ductal organization in breast carcinoma. The challenges we faced with this novel biomedical data are discussed.

**Methods:** Whole-organ and region-of-interest scans of three types of human organs were acquired at the European Synchrotron Radiation Facility (ESRF) with isotropic voxel sizes ranging from 20 µm to 0.8 µm. For the colon, voxel binning and RootPainter were employed to tackle data size to segment the ENS. For the heart, voxel-wise myocyte orientation mapping was calculated in terabyte-scale datasets with a high-performance computational framework (Cardiotensor). Breast carcinoma samples were correlated with histopathology for structure validation.

**Results:** HiP-CT revealed the large-scale organization of the ENS in the colon, enabling visualisation of the 3D structures of the ENS across the colon In TOF hearts, analysis uncovered abnormal myocardial structure and heterogeneous conduction system morphology. In breast carcinoma, HiP-CT resolved the full hierarchy of ductal structures and vascular relationships within tumour and peritumoral regions.

**Conclusions:** HiP-CT provides unprecedented, hierarchical insight into intact human organ structure, bridging the gap between histology and radiology.

**Advances in knowledge:** HiP-CT establishes a new ex vivo radiological modality capable of linking microscale pathology to whole-organ context, advancing translational research in neurogastroenterology, cardiology, and oncology

## Introduction

In 2020, a breakthrough in biological imaging was achieved with the first multi-scale phase contrast x-ray tomography and hierarchical phase-contrast tomography (HiP-CT) of entire human organs(1, 2). This is possible through X-ray phase-contrast propagation-based technique which utilizes the extremely brilliant source of the European Radiation Synchrotron Facility (ESRF). HiP-CT provides hierarchical resolution, enabling whole-organ scans at 8–25 µm voxel size and zooming into selected regions-of-interest (ROI) at voxel sizes down to 0.8 µm without the need for sectioning of the organ. Thus, HiP-CT bridges a critical gap between clinical imaging and histopathology, offering histology-like detail within the 3D structural context, without destroying the integrity of the organ.

The radiation dose required to achieve acceptable signal to noise ratio is several orders of magnitude too high for in vivo scans. As such, HiP-CT represents a new frontier in *ex vivo radiology*, a modality that, while performed post-mortem or on resected samples, holds significant translational potential to improve *in vivo* diagnostics by uncovering disease mechanisms and identifying potential therapeutic targets(1, 3). Since September 2023, the Human Organ Atlas hub (HOAHub), a worldwide collaborative effort consisting of computer scientists, biologists, physicists, and physicians, has been working on the development and application of HiP-CT to clinical challenges. This wide range of expertise is essential to improve and exploit the technique and to address key biomedical challenges. To date, the consortium has scanned a total of 263 human organs, generating 1018 individual datasets, resulting in 607TB of reconstructed data, with 297 datasets already published with open access (4, 5). The technique was developed during the COVID-19 pandemic, where it provided new insights into this disease. HiP-CT revealed arteriovenous shunting in the lungs of patients with severe COVID-19 and identified micro ischemia of the secondary lobule as a driver of fibrotic changes, supporting evidence to administer anticoagulative drugs during a time of great global and clinical uncertainty (3, 6).

The massive data generated by HiP-CT present challenges. Simply loading them for visualization requires specialised high-performance servers. Segmentation and quantification, to answer a research question or support an hypothesis, requires advance computational pipelines and substantial computing power. Through the colon, heart, and breast carcinoma case studies, we describe the technical and analytical challenges encountered when working with large HiP-CT datasets and illustrate how HiP-CT can reveal multi-scale anatomical structures that are inaccessible to conventional imaging or histology. Together, these examples provide practical insight for researchers and clinicians interested in adopting HiP-CT and highlight the unique potential of this technique to link microscale pathology with whole-organ context.

## Material, Methods and Results

### The Colon

Constipation is a frequent occurring condition, which is often well treated. However, some patients have therapy-resistant constipation (TRC), with has a strong negative impact on their health-related quality of life (7). This condition is yet not fully understood, but biological mechanisms for TRC might be found in the large scale 3D architecture of the nerves and musculature, which have been left unexplored (8). The enteric nervous system (ENS) is a network of nerves, ganglia and supportive cells that innervates the colon. These structures are microscale (50 μm diameter for nerve bundles with individual fibres and axons in the submicron range) and could only be observed through histological sectioning. Such methods forfeit the 3D orientation, and therefore the full network has been scarcely investigated(8, 9). Thus, there is no knowledge on what the 3D ENS network looks like over larger sections of the colon in either health or disease. To investigate the ENS, HiP-CT imaging was performed on an adult human colon (LADAF-2021-17, a 63 year old male. Cause of death was pancreatic cancer). An overview scan of the entire organ was acquired at 20 µm per voxel, followed by ROI scans at 4.5 and 1.4 µm voxel size in regions of interest at beamlines BM18 and BM05 of ESRF(10–14).

#### Data size management

The first challenge in analysing this data was its large size. The overview scan of the colon at 20 µm voxel size was 1 TB in size, exceeding the RAM capacity of most high-end desktop computers. Binning the data, i.e. 2×2×2 voxel fusion, reduced the data size by 8-fold, but at the expense of halving the isotropic voxel size to 40 µm. To analyse a feature, typically at least 3 voxels in its diameter are required to be able to distinguish it (15), meaning that the smallest feature that can be analysed in the binned dataset was approx. 120 µm, which was not sufficient for ENS analysis. Nerves of the myenteric plexus could be distinguished in the binned 4.5 µm scan, with a voxel size of 9 µm which allowed for identification of nerves of approximately 27 µm diameter or larger (**Fig. 1**).

**Fig 1.**
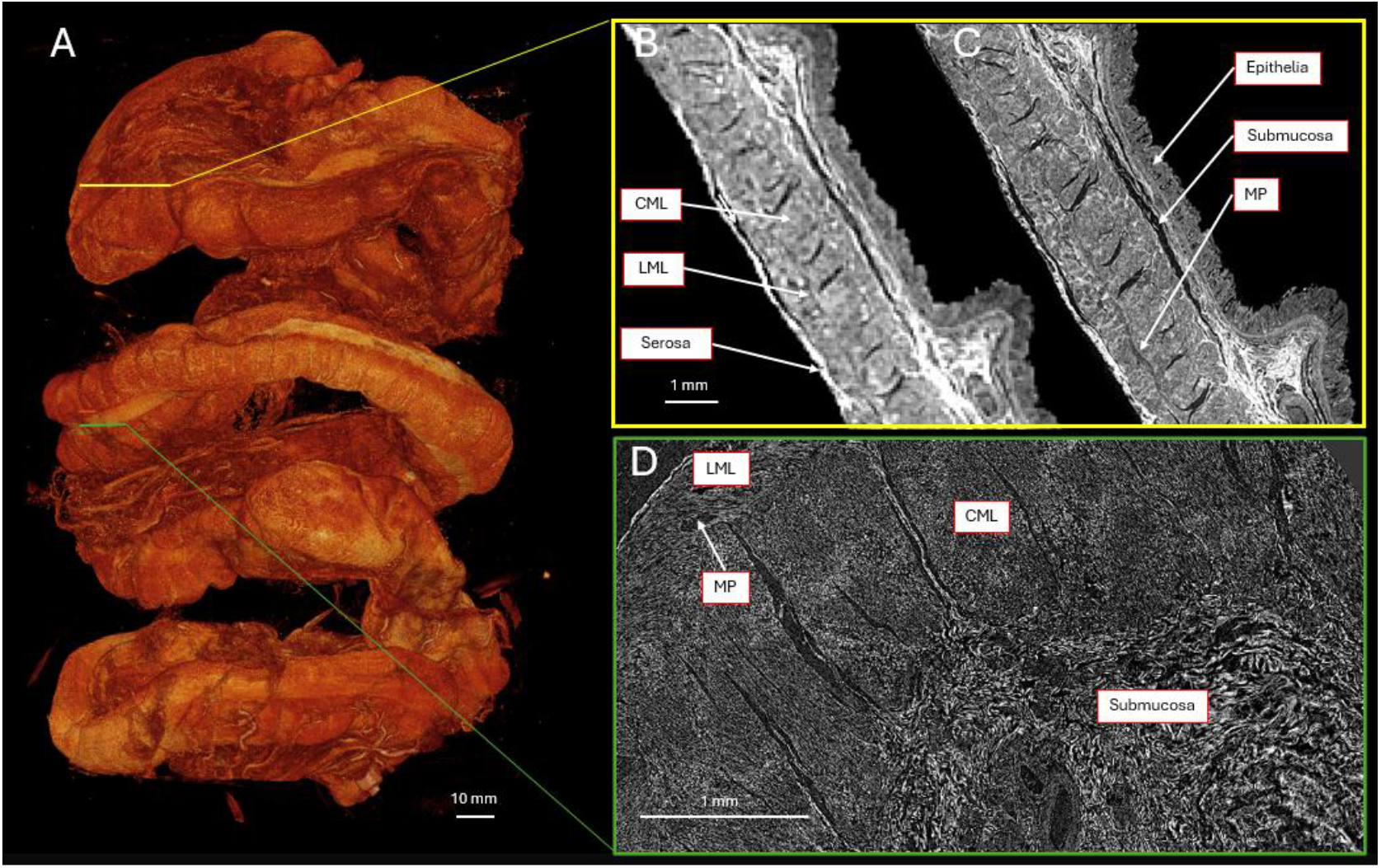
HiP-CT images of the human colon. **A**. 3D rendering of the full organ displayed at a voxel size of 40 µm. **B**. Single slice of the 40 µm whole-organ scan, showing the possibility to differentiate colon wall elements. **C**. High resolution local ROI scan shown at 9 µm voxel size, revealing more structures including the nerves of the myenteric plexus. **D**. Highest resolution ROI scan at 1.4 µm voxel size. Individual muscle fibres can be observed as well as vessels and nerves within the submucosa. **Legend** CML-Circular Muscle Layer, LML – Longitudinal Muscle Layer, MP-Myenteric Plexus

#### Segmentations and quantification

The second challenge was segmentation of the nerves for 3D visualisation and quantification, which was done on the 9 µm voxel scan. Classically this is done through manual segmentation. However, a single colon scan consists of 8000 slices and other scans in the Human Organ Atlas regularly exceed 10,000 slices. As such, it would be extremely labour-intensive and near impossible to do manual segmentation for a larger sample size. To tackle this, we employed RootPainter(16), a U-Net based machine learning tool originally developed to segment plant roots, which share morphological similarities to nerves. RootPainter requires only a small annotated training set which can be many times smaller than the original data. We had success with annotations of 100 slices, but this can vary depending on complexity and variability of the structures of interest and quality of the scans. In this training set, structures of interest are annotated and used to train a model. The predicted output of this model is shown on more slices and was corrected if necessary, allowing for interactive training of the model, and validation by an expert. With minimal computational resources, RootPainter allowed for the segmentation of the ENS over centimeters of colon **(Fig. 2)**. The program could be run on a standard laptop, however a stronger GPU allowed for faster training.

**Fig 2.**
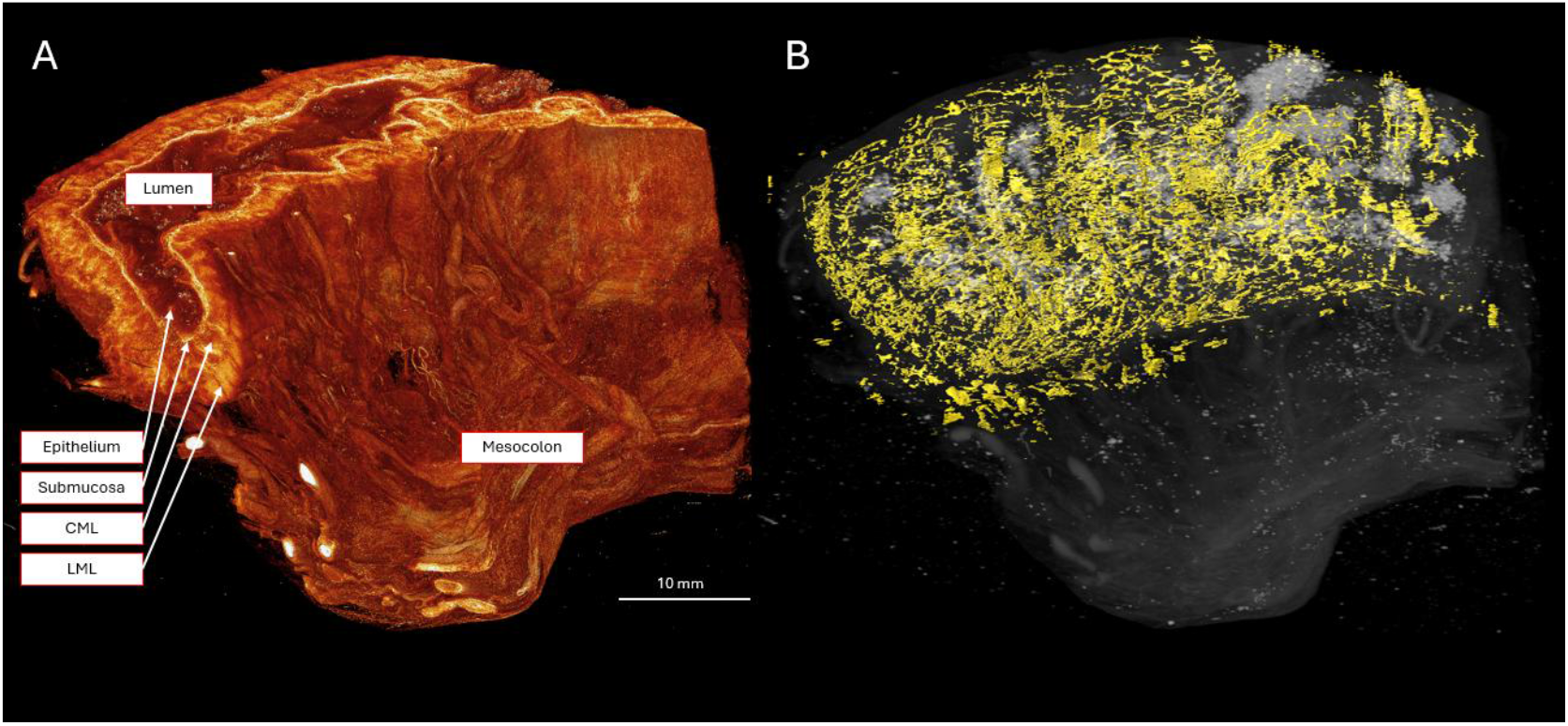
Segmentation results of the colon data. **A**. Volume rendering of the reconstructed data, allowing for differentiation of microscale structures. **B**. Segmentation of the myenteric plexus in yellow, projected over the HiP-CT data, showing its 3D architecture. **Legend** CML-Circular Muscle Layer, LML – Longitudinal Muscle Layer

### The Heart

Congenital heart disease (CHD) affects nearly 1% of live births(17). Among these, Tetralogy of Fallot (TOF) represents one of the most prevalent surgically corrected conditions. Despite decades of clinical management, the structural variability of TOF hearts and the long-term risk of arrhythmia following surgical repair are still poorly understood. A key limitation has been the inability to visualize the entire cardiac architecture, including the myocardium and conduction system, across multiple spatial scales in intact hearts. Using HiP-CT, we sought to overcome this limitation by imaging 11 whole paediatric TOF and 9 control hearts, obtained from the UCL Cardiac Archive Biobank & Birmingham Children’s Hospital, generating 3D datasets that bridge macroscopic morphology and microscopic myoarchitecture(18, 19).

#### Myomapping

Detailed analysis of myocardial architecture, especially the spatial orientation of myocytes, may enable classification of disease evolution and enhance understanding of post-operative remodelling and right ventricular dysfunction. To quantify cardiomyocyte orientation at whole-heart scale, structure tensor analysis was applied to the 9 µm HiP-CT datasets, from which helical and intrusion angles were derived. Because each dataset comprises hundreds of gigabytes, conventional desktop-based processing was not feasible. To address this, we developed a dedicated computational pipeline, Cardiotensor (20), which performs chunk-based structure tensor computation in parallel across a high-performance cluster. Parallelization and optimized memory management reduced the computation time for a full 9 µm heart from 2 days to 2 hours, while maintaining voxel-wise accuracy. The resulting vector fields were used to derive quantitative maps of helical and intrusion angles, and tractography, providing a continuous description of the myocyte aggregate orientation across the entire myocardium (**Fig. 3**).

**Fig 3.**
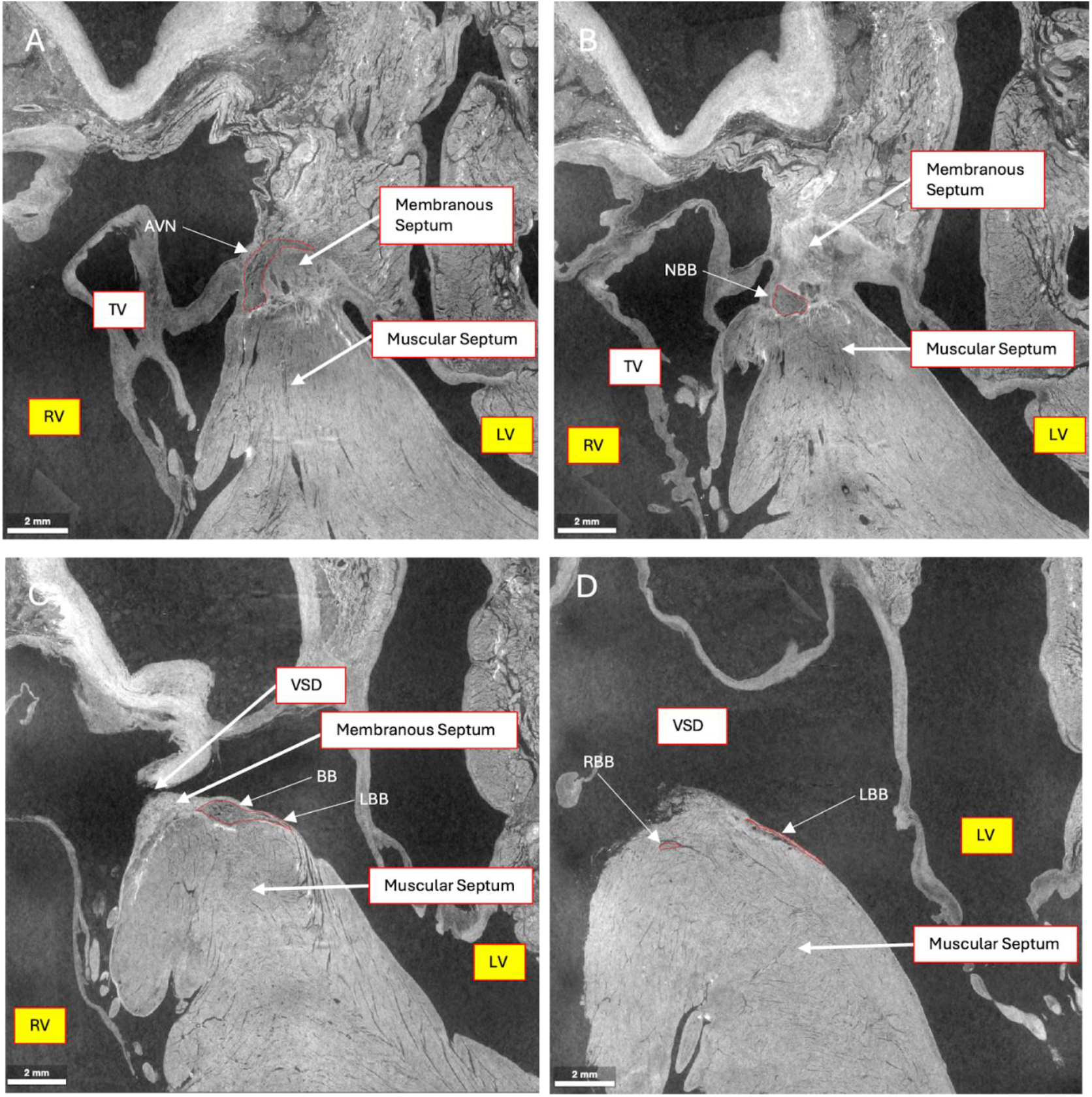
HiP-CT images of a Tetralogy of Fallot specimen scanned at 10 µm voxel size. Conduction tissue is outlined in red. AVN – atrioventricular node, TV – tricuspid valve. RV – right ventricle, LV – left ventricle, NBB – non branching bundle (of His), BB – branching bundle, LBB – left bundle branch, RBB – right bundle branch.

#### Mapping the atrioventricular conduction system

The second challenge concerned segmentation of the cardiac conduction system. Unlike purely fibrotic or vascular structures, the conduction axis is made of specialised myocytes and exhibits similar contrast to surrounding myocardium in HiP-CT datasets, albeit sheathed within fibrous components for some of its course (**Fig. 4**). Automated segmentations were unable to accurately segment these features. This was addressed with high-resolution ROI at 2 µm in targeted regions, identified from the whole-organ data, allowing segmentation of atrioventricular conduction tissue according to standard histologic criteria (Amira, Thermo Fisher Scientific, US). These local annotations in the ROI scans were then interpolated and registered back into the global volume to reconstruct the complete conduction axis in Amira (Thermo Fisher Scientific, US). This hierarchical approach preserved spatial context while ensuring anatomical accuracy.

**Fig 4.**
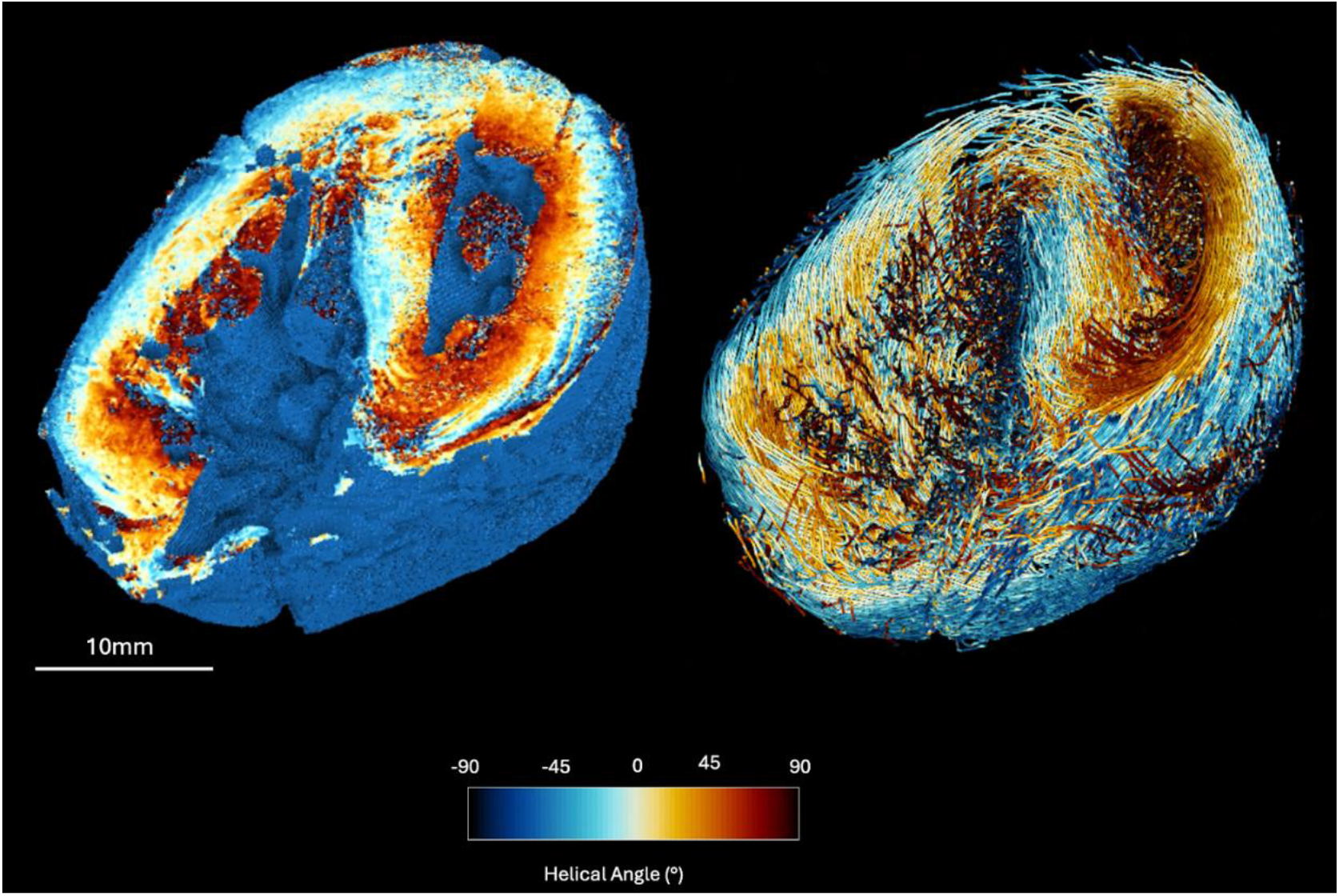
Myocyte orientation analysis in a whole-organ scan of a Tetralogy of Fallot specimen. 3D rendering of mid-ventricular slice of a ‘myomapped’ Tetralogy of Fallot specimen (left) and subsequent tractography (right). Colors represent the helical angle of myocyte aggregates.

### Breast cancer

Breast carcinoma remains the most prevalent malignancy and leading cause of cancer-related death among women worldwide (21, 22). Breast cancer typically originates from ductal epithelial cells (23, 24). Although microscopic histology provides detailed insight into ductal walls and tumour morphology, it remains inherently limited to two-dimensional sections, making histopathological work-up of cancer specimen time consuming and complex. As a result, spatial continuity across tissue volumes is lost, making it challenging to map how ducts integrate within and around tumour regions in three dimensions. This limitation can obscure essential information about tumour growth patterns, neoadjuvant therapy efficacy and resection borders.

HiP-CT imaging was performed on a full surgical sample from a 63-year-old female who underwent a partial mastectomy. The specimen contained an invasive carcinoma with a tumour diameter of 1.2 cm. The overview scan was acquired at 10 µm voxel size and 2 ROI scans were performed at 4 µm voxel size targeting the tumour margins (**Fig. 5**).

**Fig 5.**
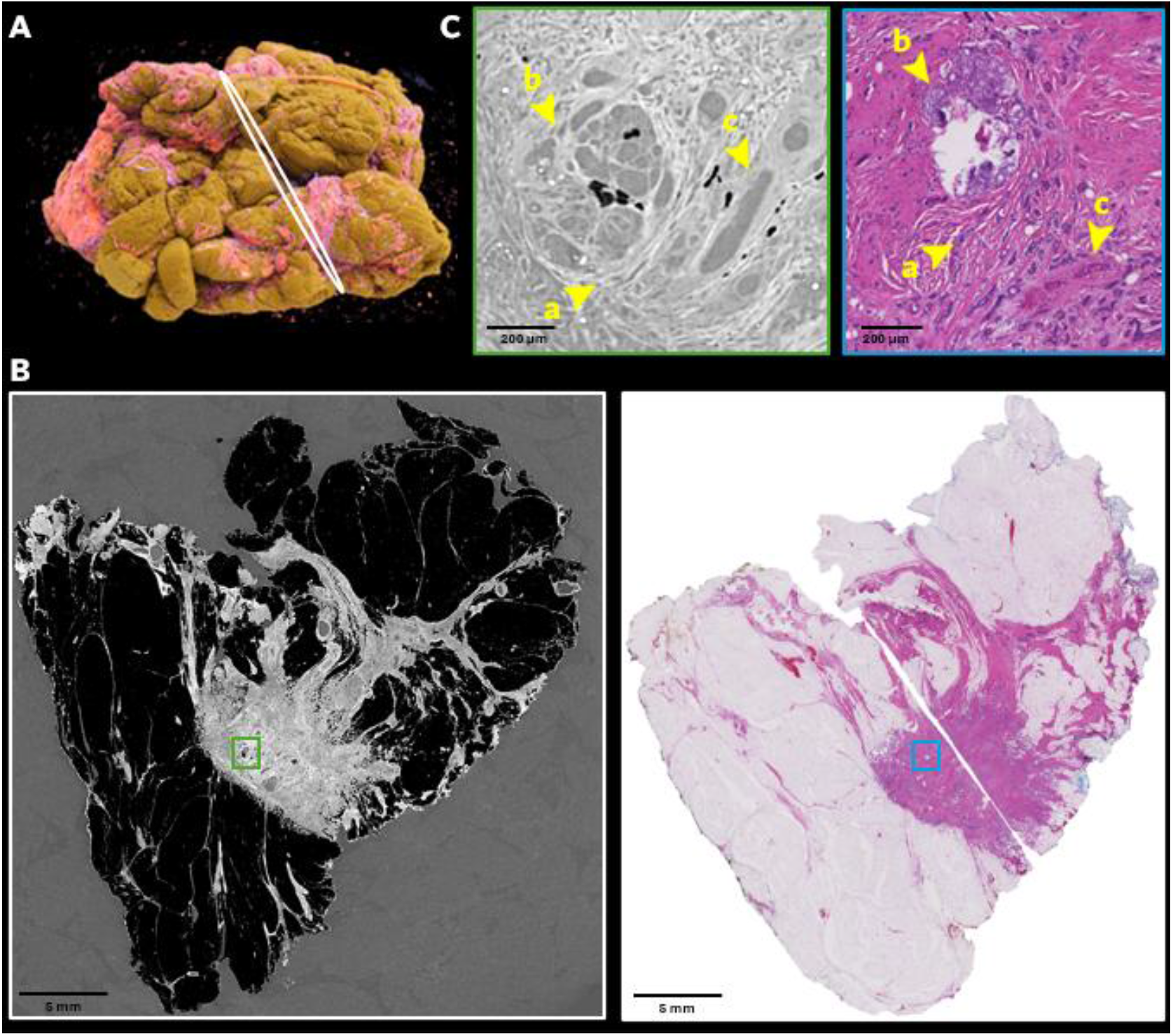
Comparison of histology and HiP-CT imaging of a human breast carcinoma of no special type (NST). **A**. Cinematic rendering of the whole breast tumor region (Cinematic Anatomy, Siemens Healthineers, Germany), with an indication of the 2D data slice (white). **B**. HiP-CT data slice of the same specimen, acquired at an isotropic voxel size of 10 µm (left) and corresponding stitched H&E-stained histological section (right). **C**. A magnified view of the HiP-CT data (at 4 µm voxel size) and histology, highlighting a representative area containing fibrosis and invasive tumor components (a), a precursor lesion (b), and associated vasculature (c).

#### Correlative imaging

Because HiP-CT is a novel modality, interpretation requires validation with established methods, especially in clinical contexts. In HiP-CT, ducts and blood vessels often exhibited similar grey-value intensities and morphological features, making them difficult to differentiate reliably. To address this limitation, corresponding histopathological sections were used as reference. Histology is typically obtained from smaller, separately oriented and processed tissue blocks that inevitably differ from the original specimen in orientation, cutting angle, rotation, and non-linear deformation during processing. The number of degrees of freedom that can occur when taking histological sections from a large sample leads makes manual registration highly challenging. Moreover, the difference in contrast mechanics between HiP-CT and histology makes automated registration a highly complex process. These two challenges are compounded by the large volumetric datasets of HiP-CT. However, features from the histological slides could be recognised in the HiP-CT data and thus an approximate slice could be easily identified. This co-registration proved essential for correct identification of ducts, arteries, and other microstructures in the HiP-CT data that could then be analysed in 3D.

## Discussion

### Biomedical challenges

We describe three biomedical challenges that were investigated within the HOAHub framework using HiP-CT imaging. These case studies highlight the potential of HiP-CT imaging for investigating microscale structures in macroscale samples in 3D orientation, a new area that may hold answers to long-existing unsolved biomedical challenges. The pipeline required to perform this research however is complex and requires multidisciplinary collaboration. Biomedical scientist and clinicians, including radiologists, pathologists and disease specialists, are needed to provide research questions, samples, interpret results, annotate the images and ultimately valorise the research into clinical solutions. Physicists and engineers are required to build the infrastructures necessary, and perform experiments. Lastly, data scientists and AI specialists have the expertise to quantify and analyse the large data that is generated. These processes however are not linear, but require close collaboration of these specialists at every step of the way to go from clinical challenge to sustainable and scalable solution.

#### The Colon

HiP-CT enabled visualization of the nerves of the ENS over large segments of colon. The TBs of data that this generates can be analysed with machine learnings tools such as RootPainter. Through this, we were able to differentiate the orientation of the ENS in 3D and improve our understanding of its macroscale network. Further research with larger samples sizes might improve our understanding of the fundamental principles behind TRC, but can be extrapolated to more diseases, ultimately shaping new research ideas that can aid these patients.

#### The Heart

Previous literature promotes a ‘three layer’ description (25, 26) of myocardial organisation within the right ventricle in TOF. Our findings however demonstrate a more gradual and continuous transition of myocyte orientation from endocardium to epicardium with the right ventricular becoming more ‘left ventricular’ in structure. The AV conduction axis, has not been described in the literature since the 1980s, or even the 1950s (27–30) and showed marked heterogeneity, particularly of the right bundle branch. We found a more ‘embryonic’ pattern evident in the shape of the atrioventricular conduction system as a whole, suggesting a lack of remodelling of the conduction system in TOF. This data could be used for the development of an updated surgical guide for VSD closure in TOF, with the aim of reducing post-operative arrhythmia.

#### Breast Cancer

HiP-CT demonstrated the capacity to resolve the 3D ductal components including the lactiferous ducts, inter- and intralobular ducts, terminal duct lobular units (TDLUs), and acini, within a single imaging modality at near histological resolution (**Fig 5)**. Such integrative 3D characterization holds promise for improving our understanding of branching complexity, tumour–stroma interactions and microenvironmental remodelling in breast carcinoma, ultimately offering new structural insights that complement and refine current histopathological and radiological assessments. Such groundwork provides a methodological framework for accurate multimodal registration of histology with synchrotron-based 3D imaging.

### Challenges in segmenting 2D and 3D

Segmentation, particularly instance segmentation of discrete anatomical entities (e.g., cardiac conduction system or glandular architecture), is an enabling technology for quantitative pathology that can provide novel morphometric landmarks and biomarkers(31). In routine 2D histopathology, field advancement is constrained less by architectural novelty of segmentation models than by the data regime. Histopathology images are extremely large and highly variable in appearance due to inter-laboratory differences; diagnostically relevant patterns span multiple magnifications, yet there are limited freely available datasets for training sufficiently robust segmentation models. These properties are unfavourable for instance segmentation, where boundary precision and object separation are critical: label noise arises from sectioning artifacts, staining variability, and inter-observer differences.

Additionally, the tiling strategies required for whole-slide images (WSIs) can decouple local texture from global context, complicating generalization across laboratories, scanners, and cohorts.

In 3D histopathology (e.g., HiP-CT), these challenges intensify because volumetric annotation is more expensive than 2D annotation and consistency constraints across volumes must be maintained. The limited experience of pathologists with 3D histological visualization introduces a novel challenge: defining histological instances volumetrically, which complicates annotation guideline development. These constraints make segmentation ground truth generation a major bottleneck.

Potential solutions include shifting to human-in-the-loop training approaches, where annotation becomes an iterative, model-guided process that reduces overall annotation burden(32). Complementary techniques - including weak supervision (using points, boxes, scribbles, or partial contours(33)), pseudo-labeling/self-training(34), and active learning(35) - can further reduce annotation requirements while enhancing coverage of rare morphologies. Segmentation foundation models, such as SAM and MedSAM derivatives, are most beneficial not as standalone pathology solutions but as proposal engines that accelerate curation by generating high-recall candidate instances from sparse prompts (points/boxes). Foundation model-assisted ground truth generation is feasible for both 2D histopathology (e.g., using QuPath) and 3D data (e.g., using 3D Slicer) via plugins also for users without significant coding experience. However, since foundation model-generated masks can be plausible but incorrect for histopathology-specific textures and domains, current best practices involve coupling them with lightweight adaptation techniques such as adapter/LoRA-style fine-tuning(36), prompt tuning, or small task heads alongside pathology-informed priors.

### Future directions

The capabilities of HiP-CT are expanding and with significantly reduced scan times achieved recently, comes an exponential increase in data volume. Working with large data is challenging and generating quantitative information for analysis from HiP-CT scans requires automated solutions, as manual analysis is already infeasible at current data volumes (37). However, computational models themselves are not the end goal, their outputs tailored to the clinical question at hand are. Because the required outputs differ per clinical question and tissue type, the extrapolation of a singular computational model is therefore limited and transfer learning might not be possible for a next project. This means that for every project, new models need to be developed and the overall timesaving capabilities of AI applications is therefore still limited. However, the basis for many of the solutions that we found for specific challenges can be used for other challenges. Cardiotensor could be used to analyse orientation of other structures, RootPainter can also work for other structures than nerves and morphological learning through co-registration with histology will form a basis for HiP-CT interpretation of all tissue.

We are now in an era, where the data from a handful of samples holds more information than can reasonably be extracted. BM18 at ESRF will shortly be technically ready to scan a whole human body at 20 µm voxel size. This will result in a reconstructed dataset of 125 TB, a size that presents even basic data storage issues. The raw data of a single scan however will be over 200 TB, and experiments spanning multiple days can even generate petabytes of raw data, from just a handful of samples. Analysing this data is extremely difficult, even with AI. This is opposite from the current medical research landscape, were we have little information per patient, but a great number of patients. This requires adaptation from our standard research rituals and move from quantitative to a qualitative research format. This provides a statistical problem, as we will be less likely to prove significant differences between groups with a small sample size. We however will be able to look more holistically at our samples, which fits current clinical developments (38). Personalized medicine is gaining more ground and we start to ask not what is a number to treat for a medicine, but why is this medicine not working for my patient (38). HiP-CT fits this development, where we can look at huge fields of views in organs, or soon even a full human body, with exceptional resolution. This means we need to start asking the right questions, ones that we might not have thought of yet, that can valorise HiP-CT data to its fullest and ultimately aid our patients.

## Ethics

All organs used in this study were obtained with ethical approval of local committees for the corresponding projects. The colon used in this study was obtained through the Laboratoire d’Anatomie des Alpes Françaises (LADAF), following full ethical approval and in accordance with French regulations for body donation and international ethical standards. Hearts were acquired from the UCL Cardiac Archive Biobank (UK Human Tissue Authority Research License 12220) and disease-free control specimens from Birmingham Children’s Hospital (BCH) (ethical approval: REC 22/PR/0906) were selected and assigned a unique ID number under UCL CAB. Transport authorization was issued by the French Ministry of Higher Education and Research N° IE-2023-2966 (UCL, London) and IE-2023-3199 (RWTH, Aachen).

## Data statement

The Human Organ Atlas hub strives towards the FAIR principles of data sharing and therefore makes data openly available at human-organ-atlas.esrf.eu. This manuscript focuses on challenges in data analysis across projects within the hub. The individual projects discussed in this manuscript are still ongoing, and their respective data will be added to the repository after their completion.

## Acknowledgements

We gratefully acknowledge the European Synchrotron Radiation Facility (ESRF) beamtimes MD1290 and MD1389 on BM05 and BM18 as sources of the data, together with all the supporting ESRF and HOAHub staff. Moreover, we sincerely thank the donors and their familie, and archieves and biobanks, for their invaluable contributions.

